## Supplementary figures and tables for "Structural analysis of 3’UTRs in insect flaviviruses reveals novel determinant of sfRNA biogenesis and provides new insights into flavivirus evolution"

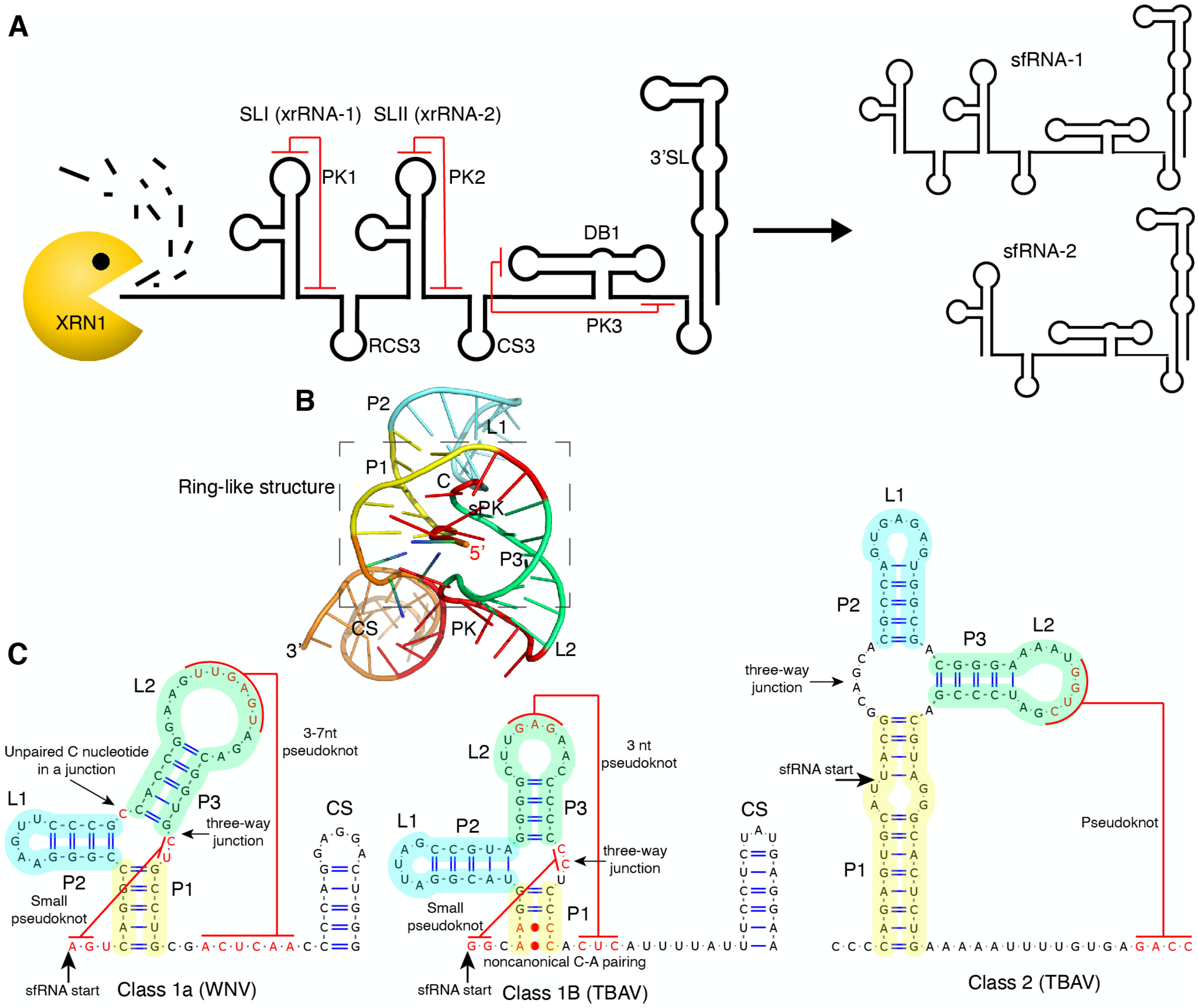


**Supplementary figure 1. Known structural determinants of sfRNA biogenesis. (A)** Schematic representation of 3'UTR organisation in mosquito borne dual-host flaviviruses (ZIKV) and biogenesis of sfRNAs via XRN1 resistance mechanism. Conserved structural elements of MBF 3'UTRs are labelled. SL – stem-loop, DB – dumbbell, PK – pseudoknot, CS3 and RCS3 – conserved small stem-loops. **(B)** Crystal structure of MVEV xrRNA (PDB ID: 4PQV)[1] shown as an example of flavivirus xrRNA tertiary structure with a 5'-end of RNA passing through ring-like element. **(C)** Secondary structure of different classes of flavivirus xrRNAs. Secondary structure of WNV xrRNA-1[2], TBAV xrRNA[3] and TBEV xrRNA-1[4] are shown as representative examples of class 1a, class 1b and class 2 xrRNAs, respectively. Functionally important structural elements are indicated and shown in colour. Colours used for highlighting match (B) to show the position of respective RNA helices and pseudoknots. In (B,C) P1-P3 – RNA helices, L1,2 – loops, CS – conserved small stem-loop.

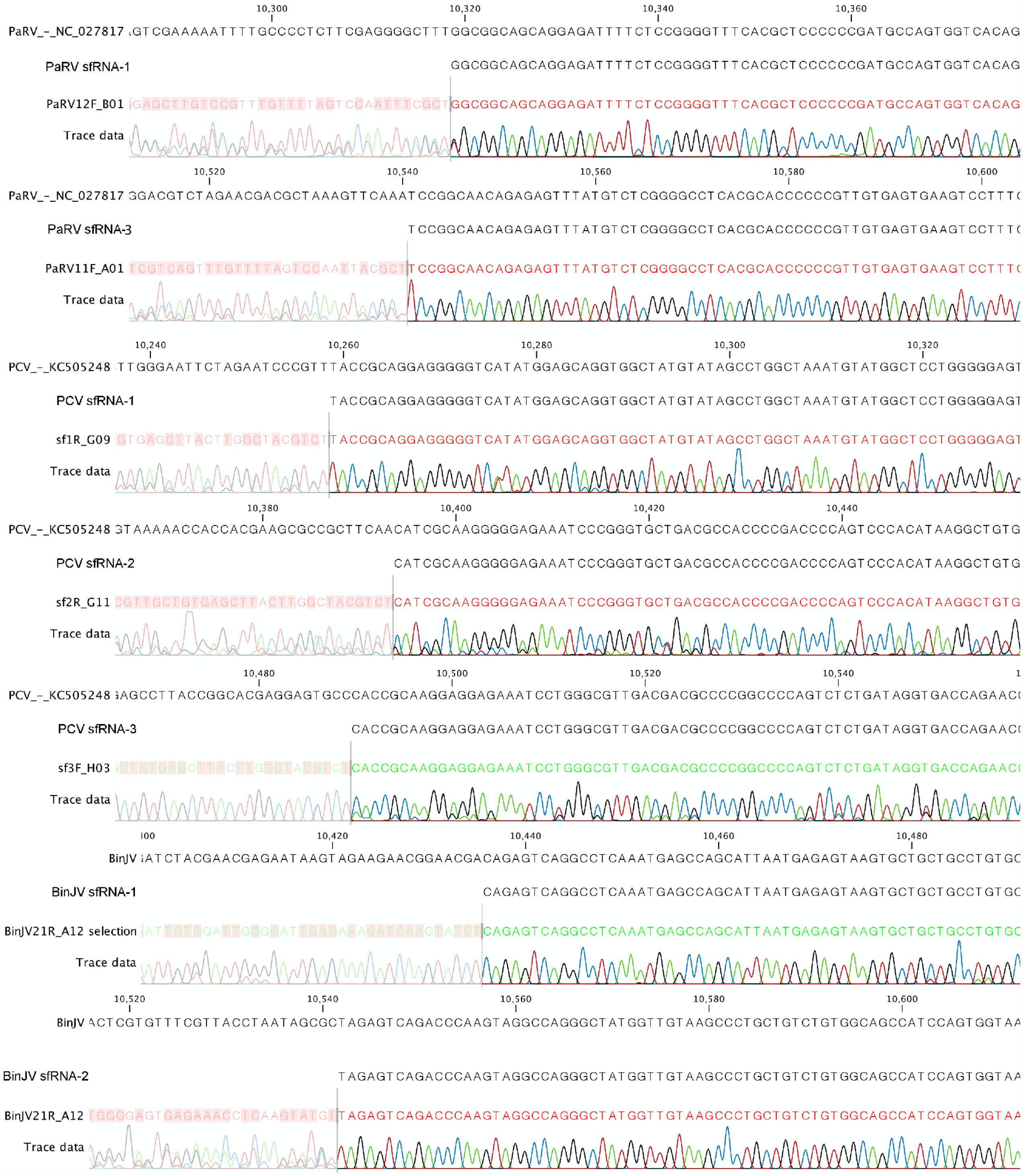


**Supplementary figure 2. Sequence alignments showing 5'-ends of ISF sfRNAs.** C6/36 cells were infected with PaRV, PCV or BinJV at MOI=1, and total RNA was isolated at 7dpi. RNA was then incubated with RNA-ligase I, which led to the circularisation of 5'-monophosphorylated uncapped RNA (sfRNA) but not capped viral genomic RNA. After ligation reaction, RNA was used for first-strand cDNA synthesis with a reverse primer designed to the 3'-end of viral UTR, followed by PCR amplification with back-to-back primers designed within the last 100nt of 3'UTRs. PCR products were gel-purified, and Sanger sequenced. The resulting sequences were aligned to genomic reference sequences, and 5'-ends of sfRNAs were identified as the first nucleotides downstream of the known 3'-terminal nucleotides in the amplicon's circularised molecules.

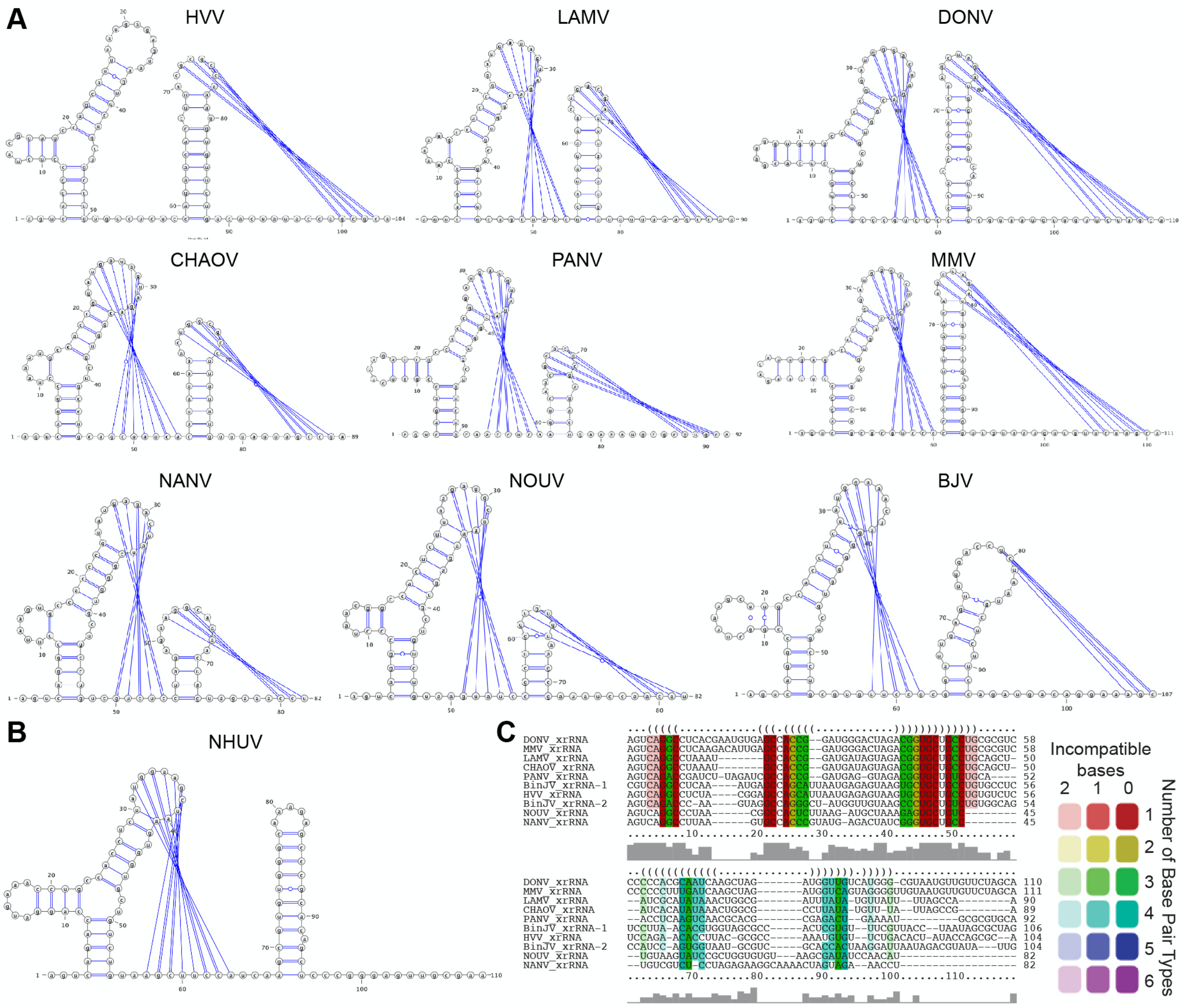


**Supplementary figure 3. conserved secondary structures of dISF xrRNAs. (A)** Predicted secondary structures of dISF xrRNAs that form novel PK. **(B)** Predicted secondary structure of Nhumirim virus. In (A, B), putative xrRNAs were located in 3'UTRs of dISFs based on sequence homology with BinJV xrRNAs. Secondary structures and pseudoknots were predicted using the IPknot web server. **(C)** Alignment of sequence and structure of dISF 3'UTR regions containing classical and novel xrRNAs. Structural constraints for alignments were specified based on experimental data (BinJV) or IP-knot prediction (other dISFs).

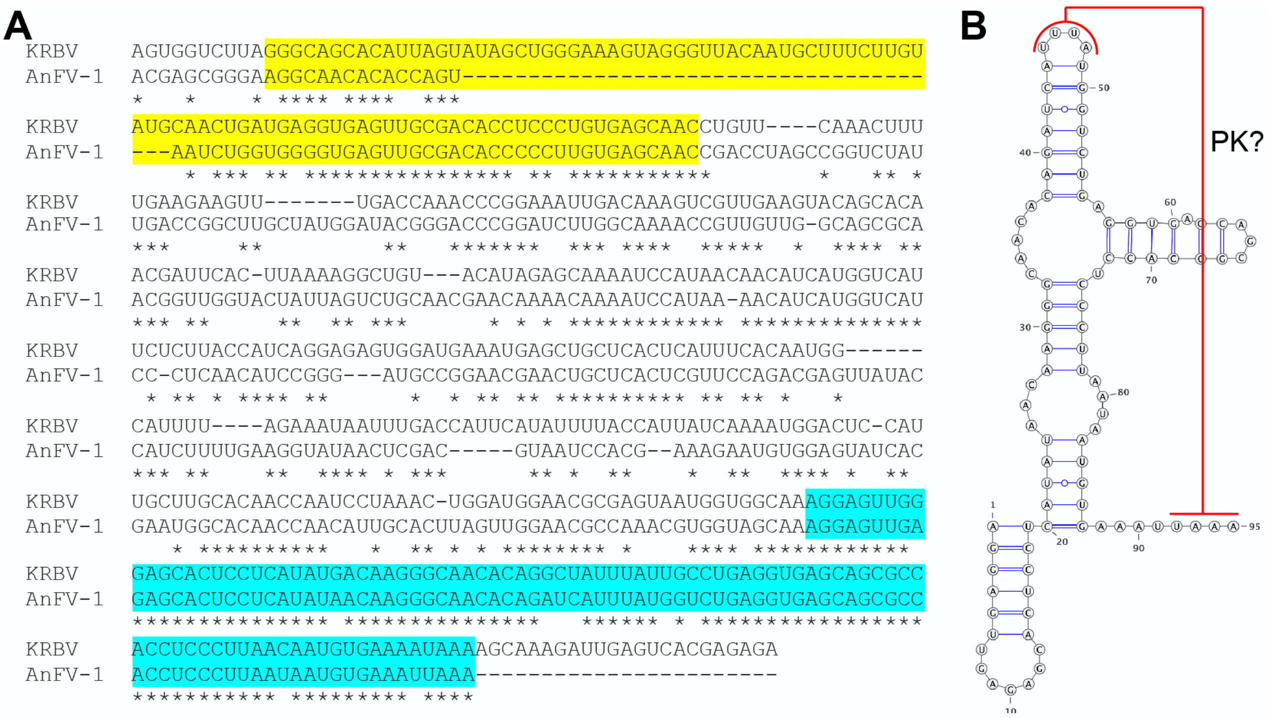


**Supplementary figure 4. Regions of homology in 3'UTRs of *Anopheles*-associated ISFs. (A)** Sequence alignment of KRBV and AnFV-1 3'UTRs. Regions corresponding to xrRNA are highlighted in yellow, and regions of 3'-terminal SL are shown in cyan. Asterisks indicate identical nucleotides. **(B)** Predicted secondary structure of AnFV-1 3'-terminal stem-loop element.

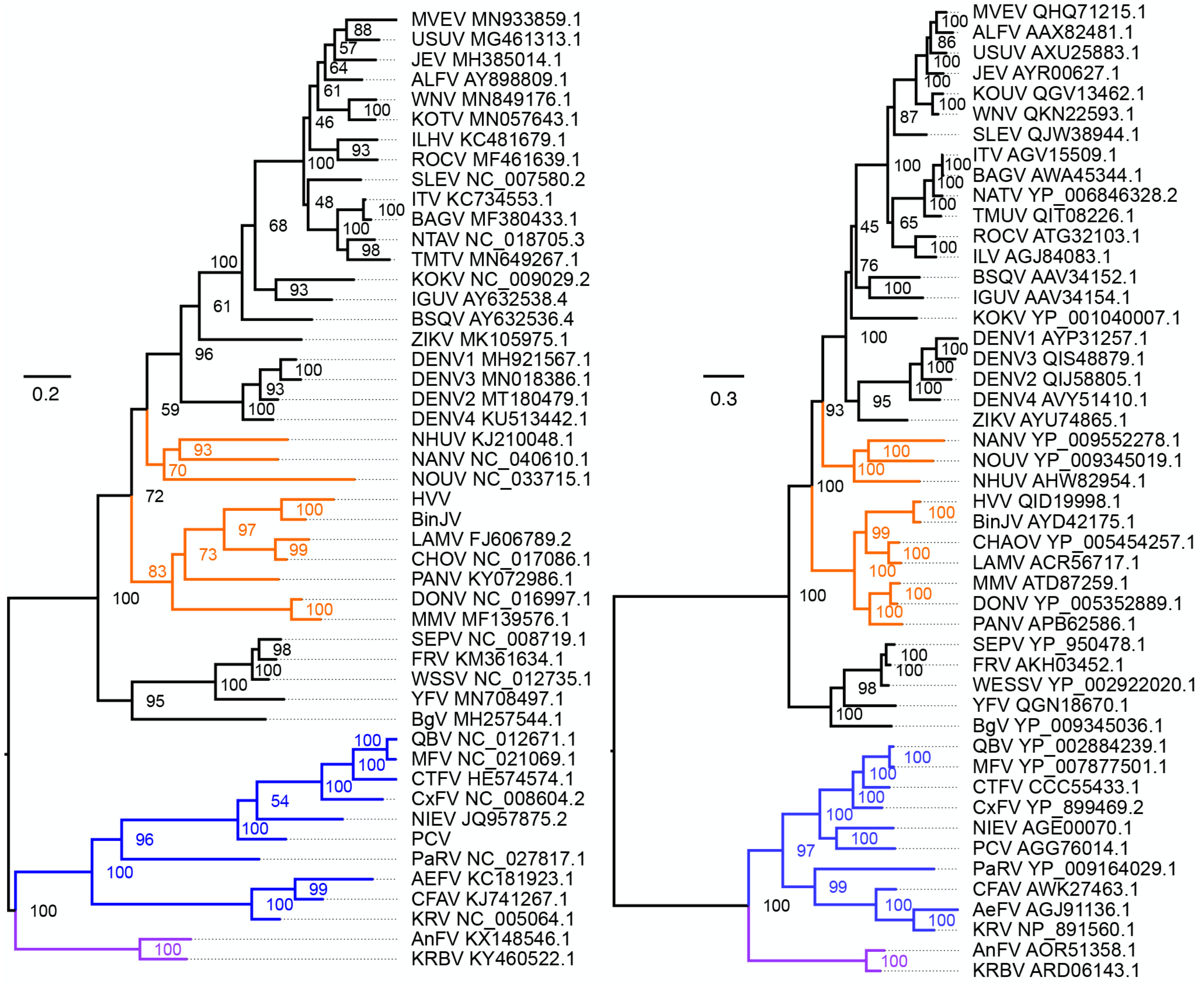


**Supplementary figure 5. Trees of phylogenetic relationships between flaviviruses built based on sequence and structure of 3'UTRs (left) and polyprotein sequence (right).** MBF clades are shown in black, dISF – in orange, cISF – in blue and *Anopheles*-associated cISFs are shown in purple. Both trees are consensus maximum-likelihood trees constructed using IQ-TREE2 and are midpoint rooted. Numbers at the nodes show bootstrap support.

**Supplementary Table 1. Virus acronym, names and Accession IDs for constructing multiple sequence alignment of flavivirus polyprotein and 3'UTRs.**

| Clade | Virus | Virus name | GenbankID 3'UTR | GenbankID |
|---|---|---|---|---|
| Anopheles cISF | AnFV | Anopheles flavivirus | KX148546.1 | AOR51358.1 |
| | KRBV | Karumba virus | KY460522.1 | ARD06143.1 |
| cISF | AeFV | Aedes flavivirus | KC181923.1 | AGJ91136.1 |
| | CFAV | Cell fusing agent virus | MH237596.1 | AWK27463.1 |
| | CTFV | Culex theileri flavivirus | HE574574.1 | CCC55433.1 |
| | CxFV | Culex flavivirus | NC_008604.2 | YP_899469.2 |
| | KRV | Kamiti River virus | NC_005064.1 | NP_891560.1 |
| | MFV | Mosquito flavivirus | NC_021069.1 | YP_007877501.1 |
| | NIEV | Nienokoue virus | JQ957875.2 | AGE00070.1 |
| | PaRV | Parramatta River virus | NC_027817.1 | YP_009164029.1 |
| | PCV | Palm Creek virus | This study | AGG76014.1 |
| | QBV | Quang Binh virus | NC_012671.1 | YP_002884239.1 |
| dISF | BinJV | Binjari virus | This study | AYD42175.1 |
| | CHAOV | Chaoyang virus | NC_017086.1 | YP_005454257.1 |
| | DONV | Donggang virus | NC_016997.1 | YP_005352889.1 |
| | HVV | Hidden valley virus | This study | QID19998.1 |
| | LAMV | Lammi virus | FJ606789.2 | ACR56717.1 |
| | MMV | Marisma mosquito virus | MF139576.1 | ATD87259.1 |
| | NANV | Nanay virus | NC_040610.1 | YP_009552278.1 |
| | NHUV | Nhumirim virus | KJ210048.1 | AHW82954.1 |
| | NOUV | Nounane virus | NC_033715.1 | YP_009345019.1 |
| | PANV | Panmunjeom flavivirus | KY072986.1 | APB62586.1 |
| MBF | ALFV | Alfuy virus | AY898809.1 | AAX82481.1 |
| | BAGV | Bagaza virus | MF380433.1 | AWA45344.1 |
| | BgV | Bamaga virus | MH257544.1 | YP_009345036.1 |
| | BSQV | Bussuquara virus | AY632536.4 | AAV34152.1 |
| | DENV1 | Dengue fever virus 1 | MH921567.1 | AYP31257.1 |
| | DENV2 | Dengue fever virus 2 | MT180479.1 | QIJ58805.1 |
| | DENV3 | Dengue fever virus 3 | MT261978.1 | QIS48879.1 |
| | DENV4 | Dengue fever virus 4 | MF004387.1 | AVY51410.1 |
| | FRV | Fitzroy River Virus | KM361634.1 | AKH03452.1 |
| | IGUV | Iguape virus | AY632538.4 | AAV34154.1 |
| | ILHV | Ilheus virus | KC481679.1 | AGJ84083.1 |
| | ITV | Israel turkey meningoencephalomyelitis virus | KC734553.1 | AGV15509.1 |
| | JEV | Japanese encephalitis virus | MH385014.1 | AYR00627.1 |
| | KOKV | Kokobera virus | NC_009029.2 | YP_001040007.1 |
| | KOUV | Koutango virus | MN057643.1 | QGV13462.1 |
| | MVEV | Murray Valley encephalitis virus | MN933859.1 | QHQ71215.1 |
| | NTAV | Ntaya virus | NC_018705.3 | YP_006846328.2 |
| | ROCV | Rocio virus | MF461639.1 | ATG32103.1 |
| | SEPV | Sepik virus | NC_008719.1 | YP_950478.1 |
| | SLEV | Saint Louis encephalitis virus | MN233334.1 | QJW38944.1 |
| | TMUV | Tembusu virus | MN649267.1 | QIT08226.1 |
| | USUV | Usutu virus | MG461313.1 | AXU25883.1 |
| | WESSV | Wesselsbron virus | NC_012735.1 | YP_002922020.1 |
| | WNV | West Nile virus | MN849176.1 | QKN22593.1 |
| | YFV | Yellow fever virus | MN708497.1 | QGN18670.1 |
| | ZIKV | Zika virus | MK105975.1 | AYU74865.1 |

**Supplementary Table 2. Oligonucleotides used in the study.**

| | |
|---|---|
| **Northern Blotting probes** | |
| PaRV NB Probe | AGCGTAATTGGACTAAAACAAACTG |
| PCV NB probe | AGACGTAGCCAAGTAAGCTCACAG |
| KRBV NB Probe | GTTAAGGGAGGTGGCGCTGCTCACC |
| BinJV NB Probe | AGATACTTGATGTTTCTCAATC |
| HVV NB Probe | ACTTGATGTTTCTCAATTCCCAATCCAC |
| **sfRNA Sequencing Primers** | |
| KRBV_Lig_Seq_F | GCGAGTAATGGTGGCAAAGGAGTTGGG |
| KRBV_Lig_seq_F | GTTCCATCCAGTTTAGGATTGGTTGTGCAAGC |
| KRBV_Lig_Seq_RT | TCTCTCGTGACTCAATCTTTGCTTTTATTTTCACATTGTTAAGGG |
| PCV_Lig_Seq_F | GGGCTTAGCCCAAGGTGAGTGACGA |
| PCV_Lig_Seq_R | TGTGTGCCCTCACCATTTCTGGTGATC |
| PCV_Lig_Seq_RT | AGACGTAGCCAAGTAAGCTCACAGCAACG |
| PaRV_Lig_Seq_F | CCCGGTTGTGAAAACGATTGCGACTAGAA |
| PaRV_Lig_Seq_R | GGGTGACGCTACTCACCCTAGTTCGTT |
| PaRV_Lig_Seq_RT | AGCGTAATTGGACTAAAACAAACTGACGAGCTCC |
| BinJV-Lig_Seq_RT | AGATACTTGATGTTTCTCAATCCCCAATCCACAATTATTCGG |
| BinJV_Lig_Seq_F | CGACACCTGGGAAAGACCGGAGATACC |
| BinJV_Lig_Seq_R | ATATGATGCTCTACATTTTGGAGGGGGTTTCCTCT |
| **Primers for generation of IVT template for XRN-1 siRNA** | |
| T7-Aedes_XRN1-1s-F | TCGCAGtaatacgactcactatagggCGAGTTGCCGAAGGAGCCGT |
| T7-Aedes_XRN1-1s-R | GCGGAACATTGGGCGGATGC |
| T7-Aedes_XRN1-1a-F | CGAGTTGCCGAAGGAGCCGT |
| T7-Aedes_XRN1-1a-R | TCGCAGtaatacgactcactatagggGCGGAACATTGGGCGGATGC |
| T7-Aedes_XRN1-2s-F | TCGCAGtaatacgactcactatagggGCCCCATCTAACCGAGGCCA |
| T7-Aedes_XRN1-2s-R | CCGATCGTCTCGCTCCTCGC |
| T7-Aedes_XRN1-2a-F | GCCCCATCTAACCGAGGCCA |
| T7-Aedes_XRN1-2a-R | TCGCAGtaatacgactcactatagggCGAGTTGCCGAAGGAGCCGT |
| **3'UTR cloning primers** | |
| PaRV-3UTR T7 F | TAATACGACTCACTATAGGGgaaaccatcttttccaaattag |
| PaRV-3UTR T7 R | AGCGTAATTGGACTAAAACAAACTGACGAGC |
| PCV-3UTR T7 F | TAATACGACTCACTATAGGGgaaaaatccttgagcaggag |
| PCV-3UTR T7 R | AGACGTAGCCAAGTAAGCTCACAGCAACGC |
| BinjV-3UTR T7 F | TAATACGACTCACTATAGGGggatctacgaacgagaataag |
| BinjV-3UTR T7 R | AGATACTTGATGTTTCTCAATCCCCAATCCACAATT |
| KRBV-3UTR T7 F | TAATACGACTCACTATAGGGccaaattacgacgtgtcttgg |
| KRBV-3UTR T7 R | TCTCTCGTGACTCAATCTTTGCTTTTATTTTCACATTGTTAAGG |
| **SHAPE Primers** | |
| PaRV_233-FAM | /56-FAM/CTAGACGTCCTTCGAAACCAGTTCG |
| PCV_243-FAM | /56-FAM/CGTCACAGCCTTATGTGGGACTGGG |
| BinJV_379-FAM | /56-FAM/TGTGCCTTGTGGATTGAGTGCTGTG |
| BinJV_200-FAM | /56-FAM/CGACGCATTACCACTGGATGGCUG |
| KRBV_379-FAM | /56-FAM/AGTGCTCCCAACTCCTTTGCCACC |
| KRBV_447-FAM | /56-FAM/CATTGTTAAGGGAGGTGGCGCTG |
| HVV_187-FAM | /56-FAM/CGGGCGCCATTCCACTATGAGGTTG |
| PaRV_233-HEX | /5HEX/CTAGACGTCCTTCGAAACCAGTTCG |
| PCV_243-HEX | /5HEX/CGTCACAGCCTTATGTGGGACTGGG |
| BinJV_379-HEX | /5HEX/TGTGCCTTGTGGATTGAGTGCTGTG |
| BinJV_200-HEX | /5HEX/CGACGCATTACCACTGGATGGCUG |
| KRBV_379-HEX | /5HEX/AGTGCTCCCAACTCCTTTGCCACC |
| KRBV_447-HEX | /5HEX/CATTGTTAAGGGAGGTGGCGCTG |
| HVV_187-HEX | /5HEX/CGGGCGCCATTCCACTATGAGGTTG |
| **3'UTR sequencing Primers** | |
| PaRV_3UTR_Seq F | AGAGACTTGGAGTTGTGGCG |
| PaRV_3UTR_Seq R | AGCGTAATTGGACTAAAACAAACTG |
| PCV_3UTR_Seq_F | AGTGTGACGATGTAGGCCCG |
| PCV_3UTR_Seq_R | AGACGTAGCCAAGTAAGCTCACAGC |
| **Mutagenesis Primers** | |
| PCV_PK1'F | CTGGGGGAGTctgGCCCCTCCGG |
| PCV_PK1'R | GAGCCATACATTTAGCCAGG |
| PCV_PK2'_F | CCCGGGTGCTctgGCCACCCCGA |
| PCV_PK2'_R | ATTTCTCCCCCTTGCGATGTTG |
| PCV_PK3'_F | CCTGGGCGTTctgGACGCCCCGG |
| PCV_PK3'_R | ATTTCTCCTCCTTGCGGTGG |
| PaRV_PK1'_F | TCCGGGGTTTgtgGCTCCCCCCG |
| PaRV_PK1'_R | GAAAATCTCCTGCTGCCG |
| PaRV_PK2'_F | CTCGGGGATTgtgGCTCCCCCCAT |
| PaRV_PK2'_R | ACAAGCTCTGCTGCCATA |
| PaRV_PK3'_F | CTCGGGGCCTgtgGCACCCCCCG |
| PaRV_PK3'_R | ACATAAACTCTCTGTTGCCGGATTTG |
| KRBV_PK1'_F | GATGAGGTGAcaacgGACACCTCCCTGTGAGC |
| KRBV_PK1'_R | AGTTGCATACAAGAAAGC |
| BinJV PK1' F | CAGCATTAATCTCTCTAAGTGCTGCTGCCTGTG |

| | |
|---|---|
| BinjJV PK1' R | GCTCATTTGAGGCCTGAC |
| BinJV PK2' F | CCAGGGCTATCCAACTAAGCCCTGCTG |
| BinJV_PK3' F | GGAGCAGCAAcgagGAGCTGCATCACCCAC |
| BinJV_PK3' R | TCACCAGTCCCTCCCAAT |
| BinJV PK3' R | CCTCACCAGTCCCTCCCA |
| BinJV nPK1' F | AACACGTGGTTCGCCCACTCGTGTTTCG |
| BinJV nPK1' R | AAGGAGAGGCACAGGCAG |
| BinJV_nPK2'F | CAGTGGTAATCGCAGGCACCACTAAGGATTAATAGACG |
| BinJV_nPK2'R | GATGGCTGCCACAGACAG |
| **CPER primers** | |
| ParV_linkerF | CAGTTTGTTTTAGTCCAATTACGCTGGGTCGGCATGGCATCTCCAC |
| ParV_linkerR | AACCACGGGTTAACTTTTAAAAACTGTTTACCAGATCGTTGCGGGC |
| ParV_frag1F | AGTTTTTAAAAGTTAACCCGTGGTTTTACC |
| ParV_frag1R | AACCGGTTTGTTCTCCCTAGTC |
| ParV_frag2F | GACTAGGGAGAACAAACCGGTT |
| ParV_frag2R | GTATGCCCTCATTACTAGCCCC |
| ParV_frag3F | GGGGCTAGTAATGAGGGCATAC |
| ParV_frag3R | CCTTCTGGCTTCCAACCATACT |
| ParV_frag4F | AGTATGGTTGGAAGCCAGAAGG |
| ParV_frg4R | AGCGTAATTGGACTAAAACAAACTGA |
| PCV_linker_F | GTGAGCTTACTTGGCTACGTCTGGGTCGGCATGGCATCTCCAC |
| PCV_linker_R | ACTAACGCAAAAGTTTTTTAAAACTGTTTACCAGATCGTTGCGGGC |
| PCV_frag1_F | AGTTTTAAAAAACTTTTGCGTTAGT |
| PCV_frag1_R | ACTCCTCTCGAACTCTCCATCA |
| PCV_frag2_F | TGATGGAGAGTTCGAGAGGAGT |
| PCV_frag2_R | CTCCCAAAGTCACCCCGATAAA |
| PCV_frag3_F | TTTATCGGGGTGACTTTGGGAG |
| PCV_frag3_R | TAACGTCAGTGGATGGAAGTGG |
| PCV_frag4_F | CCACTTCCATCCACTGACGTTA |
| PCV_frag4_R | AGACGTAGCCAAGTAAGCTCACAG |
| BinJV_5'-E_F | AACGATCTGGTAAACAGTATATTTTGCGTG |
| BinJV_5'-E_R | TCCTATTTCCGATAGGGCACCCACGGTCAC |
| BinJV_1-2B_F | GTGACCGTGGGTGCCCTATCGGAAATAGGA |
| BinJV_1-2B_R | CCACAACACAGTCCCCCGCTTGTTTGATTT |
| BinJV_3-4B_F | AAATCAAACAAGCGGGGGACTGTGTTGTGG |
| BinJV_3-4B_R | GGTGGCCTGTAATCCCCTCCTAGGAACTCC |
| BinJV_NS5-3'_F | GGAGTTCCTAGGAGGGGATTACAGGCCACC |
| BinJV_NS5-3'Junc_R | CTCGTTCGTAGATCCTTAGATCACATTGCC |
| BinJV_NS5-3'Junc_F | GGCAATGTGATCTAAGGATCTACGAACGAG |
| BinJV_NS5-3'_R | TGCCATGCCGACCCAGATACTTGATGTTTC |
| BinJV3'UTR-linker | TGGATTGGGGATTGAGAAACATCAAGTATCTGGGTCGGCATGGCATCTCCACCTCCTCGC |
| Linker-BinJV5'UTR | GTGTTTTGAAACGCACACGCAAAATATACTGTTTACCAGATCGTTGCGGGCTGTATTTATAGGC |
